# My friend MIROSLAV: A Hackable Open-Source Hardware and Software Platform for High-Throughput Rodent Activity Monitoring in the Home Cage

**DOI:** 10.1101/2024.06.25.600592

**Authors:** Davor Virag, Jan Homolak, Ivan Kodvanj, Ana-Marija Vargantolić, Ana Babić Perhoč, Patrik Meglić, Petra Šoštarić Mužić, Ana Knezović, Jelena Osmanović Barilar, Mario Cifrek, Vladimir Trkulja, Melita Šalković-Petrišić

## Abstract

While conventional behavioural tests offer valuable insight into rodent behaviour in specific paradigms, rodents spend most of their time in a safe, undisturbed environment – their home cage. However, home cage monitoring (HCM) is often impractical at scale over prolonged periods of time without significant loss of data.

To achieve this, we developed MIROSLAV, the Multicage InfraRed Open Source Locomotor Activity eValuator, a home cage activity and environmental monitoring device designed to be open, adaptable, and robust. Its transparent and modular design allows the device to be tailored to varying experimental requirements and environmental conditions, while its wireless operation and multiple redundancies minimise loss of data and animal disturbance. Data quality is maintained by a modular software workflow for data preparation (Prepare-a-SLAV) and cleaning (TidySLAV), followed by exploratory (MIROSine – MIRO The Explorer) and statistical (MIROSine – StatistiSLAV) analysis of circadian periodicity.

Here, using MIROSLAV, we demonstrate circadian dysrhythmia and a disrupted response to regular stimuli (e.g. behavioural testing, bedding change) in a rat model of sporadic Alzheimer’s disease, showcasing MIROSLAV’s utility in a typical animal study of disease.

In accordance with the 3Rs, every rodent study and laboratory performing them can benefit from HCM, provided the existence of transparent and customisable tools which allow robust deployment to tens or hundreds of cages in various conditions. MIROSLAV provides this opportunity to researchers in a cost-effective manner.

## Introduction

Circadian rhythms are present in organisms from bacteria to humans, from the molecular to the behavioural level (Merrow, 2023). Disruption of the circadian rhythm of locomotor activity has been recognised as a feature, risk factor, and possible etiopathogenetic component in a number of pathological conditions prevalent in the human population (e.g. Alzheimer’s disease (Homolak et al., 2018)). As such, it is a subject of growing interest in biomedical research, particularly in animal models of disease.

Circadian measurement of locomotor activity in laboratory rodents, however, requires a different approach than conventional behavioural tests, and is commonly performed using continuously-operating, non-invasive home cage monitoring (HCM) devices. This type of monitoring greatly increases the amount of data obtained from the same set of animals and furthers adherence to the 3Rs principle (Baran et al., 2021). In recent years, newly developed HCM systems have rapidly increased in number, particularly custom, open-source solutions (Kahnau et al., 2023).

Today, commercial tools for behavioural analysis are developed as proprietary, closed source hardware and software solutions. Proprietary assemblies and undocumented hardware communication protocols make it nearly impossible for researchers to extend a single device according to ever-changing experimental requirements. Before researchers are given the acquired data, it is automatically pre-processed and cleaned through closed source algorithms which differ across devices and manufacturers, in direct opposition to reproducibility as a fundamental scientific principle. Raw data can sometimes be obtained in an unstandardised format via explicit request to the manufacturer.

To avoid these limitations, researchers have started developing modular open-source solutions that can be customised to the requirements of novel experiments, and quickly adapted to handle unforeseen circumstances inherent to animal research, without needing to rely on a lengthy process of external testing and support. The development and publication of methods extensively documented and fully replicable in accordance with academic standards invites wider collaboration towards the uptake and further development of these novel solutions (Oellermann et al., 2022).

Inspired by previous methods built in accordance with this movement (e.g. Rodent Activity Detector – RAD (Matikainen-Ankney et al., 2019), Continuous Open Mouse Phenotyping of Activity and Sleep Status – COMPASS (Brown et al., 2016)), we propose MIROSLAV, the Multicage InfraRed Open Source Locomotor Activity eValuator, as a complete HCM hardware and software toolkit built with three main goals in mind:

1. Long-term high-throughput monitoring of hundreds of cages at a time,
2. Robustness and redundancy to prevent data loss,
3. Extensive modularity to support a wide array of experimental contexts.

This paper will first describe the MIROSLAV method, followed by a proof-of-concept study demonstrating MIROSLAV’s utility.

## Materials and Methods

### The Multicage InfraRed Open Source Locomotor Activity eValuator: MIROSLAV

MIROSLAV is an Arduino-based platform for continuous, non-invasive monitoring of home-cage locomotor activity in laboratory rodents. It consists of multiple cage-mounted sensor arrays shown as Element A of Figure 1, which register home cage activity. Cage-mounted sensor arrays are connected to the main unit’s input serialisation stack, shown as Element B of Figure 1, which feeds the readings to the main unit’s mainboard, shown as Element C of Figure 1. The mainboard timestamps the data and transmits it to an off-site computer via Wi-Fi. Data acquisition is performed with the Record-a-SLAV Python tool.

**Figure 1.**
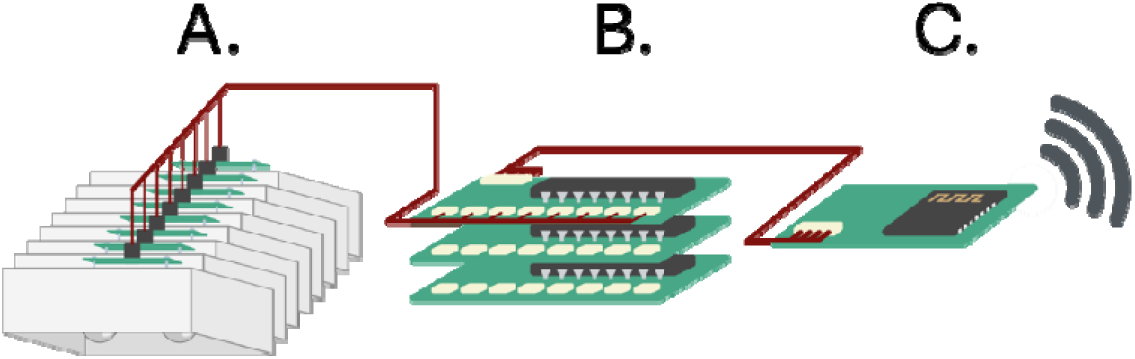
Illustration of the MIROSLAV Device and Its Components. *Note.* Element A: Cage sensor arrays. Element B: Main unit – Up to 64 input serialisation boards. Element C: Main unit – Mainboard.

The MIROSLAV hardware repository contains circuit board designs (produced in KiCad (KiCad Development Team, n.d.), with respective Gerber files, allowing customisation and quick production via online services) (Virag, 2024a), whereas the MIROSLAV firmware repository contains MIROSLAVino Arduino firmware and Record-a-SLAV acquisition software (Virag, 2024c). The MIROSLAV analysis repository contains a software toolkit for data preparation, cleaning, and analysis (Virag, 2024b).

### Cage-mounted sensor array

The home cage-mounted sensor array, shown in Figure 2, consists of two passive infrared (PIR) modules which register the rodent’s locomotor activity.

**Figure 2.**
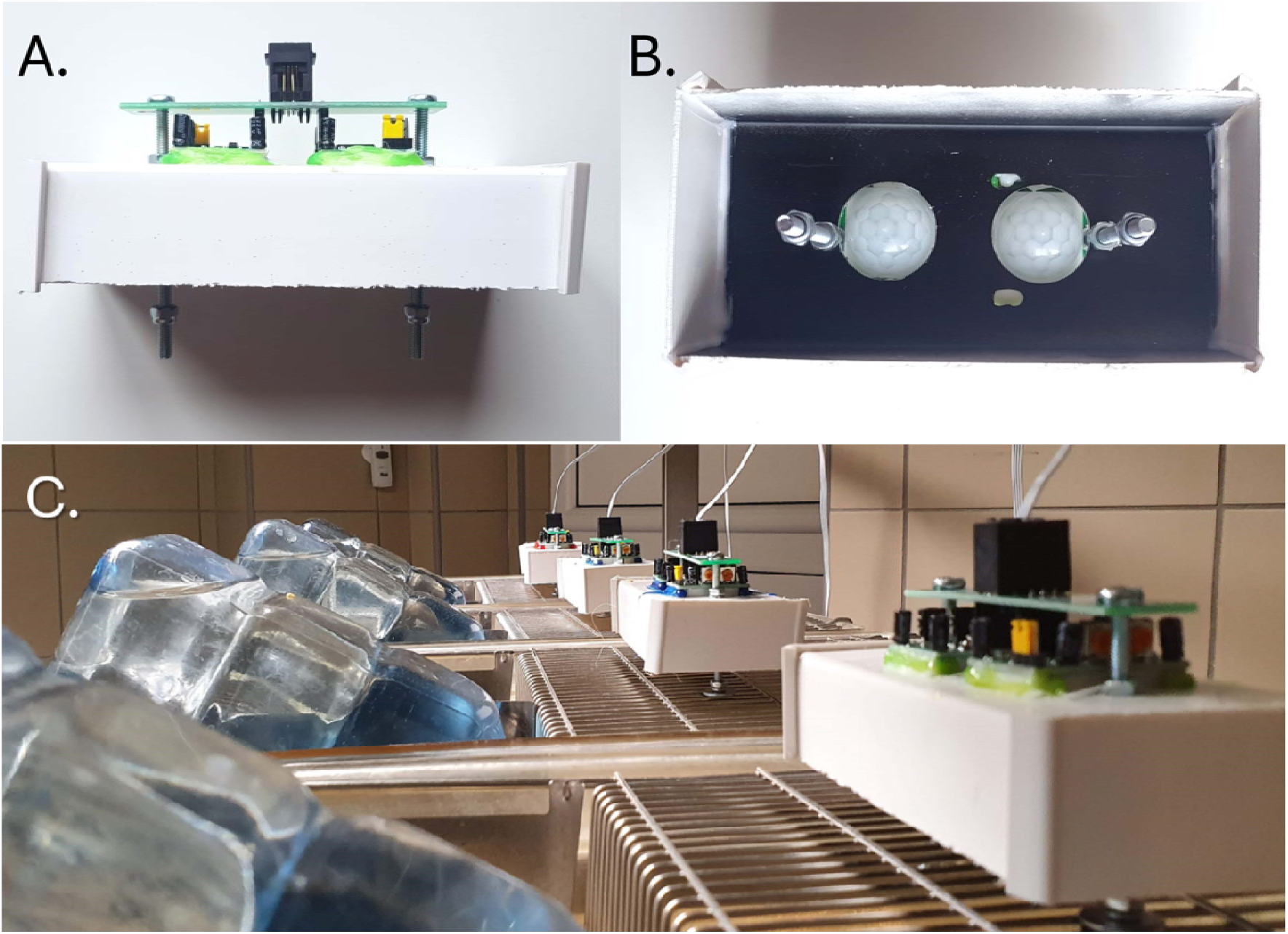
Cage-mounted Sensor Arrays in a Typical MIROSLAV Setup. *Note.* Panel A: Side view of a sensor array. Panel B: Bottom view of a sensor array, revealing Fresnel lenses of the two sensor modules. Panel C: Cage-mounted sensor arrays in operation.

The array screws onto the cage lid and its sensors are partially encased with a modified PVC cable raceway to prevent IR interference from outside the cage. On top of the assembly lies a custom printed circuit board (PCB) which provides a single easily detachable connection for power and signal output to the main unit by way of an RJ9 socket.

The main sensor component is a pair of photovoltaic cells sensitive to infrared light, paired with a segmented Fresnel lens, shown in Figure 2B, which alternately focuses segments of the field of view to each of the cells. Movement of an IR-emitting body (like that of a warm-blooded animal) in the sensor’s field of view therefore differentially excites the cells. The module’s onboard BISS0001 integrated circuit (Adafruit Industries, 2006) continuously calculates the difference between the signal of the two cells. When this difference crosses a threshold, the sensor’s output flips from a logical 0 to a logical 1, signifying movement. This results in two binary values being transmitted from each cage, one per sensor module.

Power and data transmission to and from the sensor array is done with a single 4-wire cable affixed to the rack. On the main unit side, this cable splits into two JST connectors – one for power, and one for data transmission. On the cage side, the cable converges into an RJ9 plug (similar to Ethernet and telephone landline connectors) which maintains a reliable connection and enables zero pull-out and insertion force operation once the locking pin is pressed.

### Main unit

#### Input Serialisation Stack

The input serialisation stack, shown in Figure 3A, consists of one or multiple input boards. Up to 8 individual boards may be stacked together to expand a MIROSLAV’s capacity for data input up to 64 cages (128 individual sensor modules).

**Figure 3.**
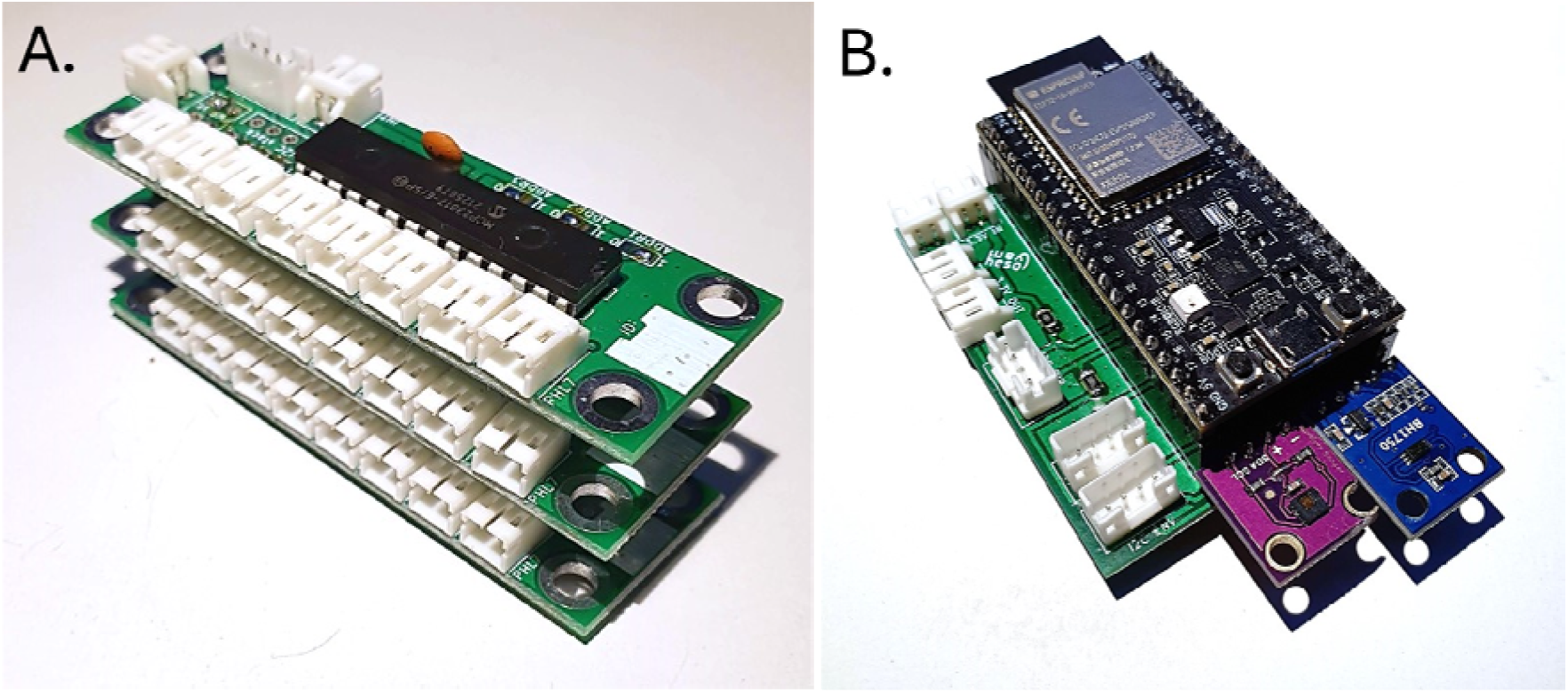
The MIROSLAV Device Main Unit. *Note.* The main unit consists of two distinct parts. Panel A: An input serialisation stack containing three input boards. Panel B: The mainboard, containing the microcontroller board and environmental conditions sensors.

Each input board consists of 8 cage input connectors, connectors for power and mainboard communication, and a stacking connector. When stacked, only the topmost board needs to be powered and connected to the mainboard. Other boards will receive power and have access to the data transmission bus towards the mainboard through the stacking connector. Thus, the entire stack can be extended to the appropriate number of inputs while acting as one unit with a single power and data connection.

#### Mainboard

The mainboard, shown in Figure 3B, is based around the Wi-Fi-enabled ESP32-S2 microcontroller compatible with the Arduino software framework. The mainboard polls the input serialisation stack for sensor data and transmits it wirelessly over the MQTT protocol to one or more computers dedicated to data logging.

Aside from the cage sensor data received from the input serialisation stack, the mainboard also records environmental data. Onboard sensors measure ambient light level, temperature, and humidity, and a PIR sensor connected to a dedicated socket monitors the habitat for personnel entrances. An example of environmental data gathered with a MIROSLAV prototype, parsed, and plotted using EnviroSLAV is shown in Figure 4. EnviroSLAV is available in the MIROSLAV analysis toolkit repository (Virag, 2024b)

**Figure 4.**
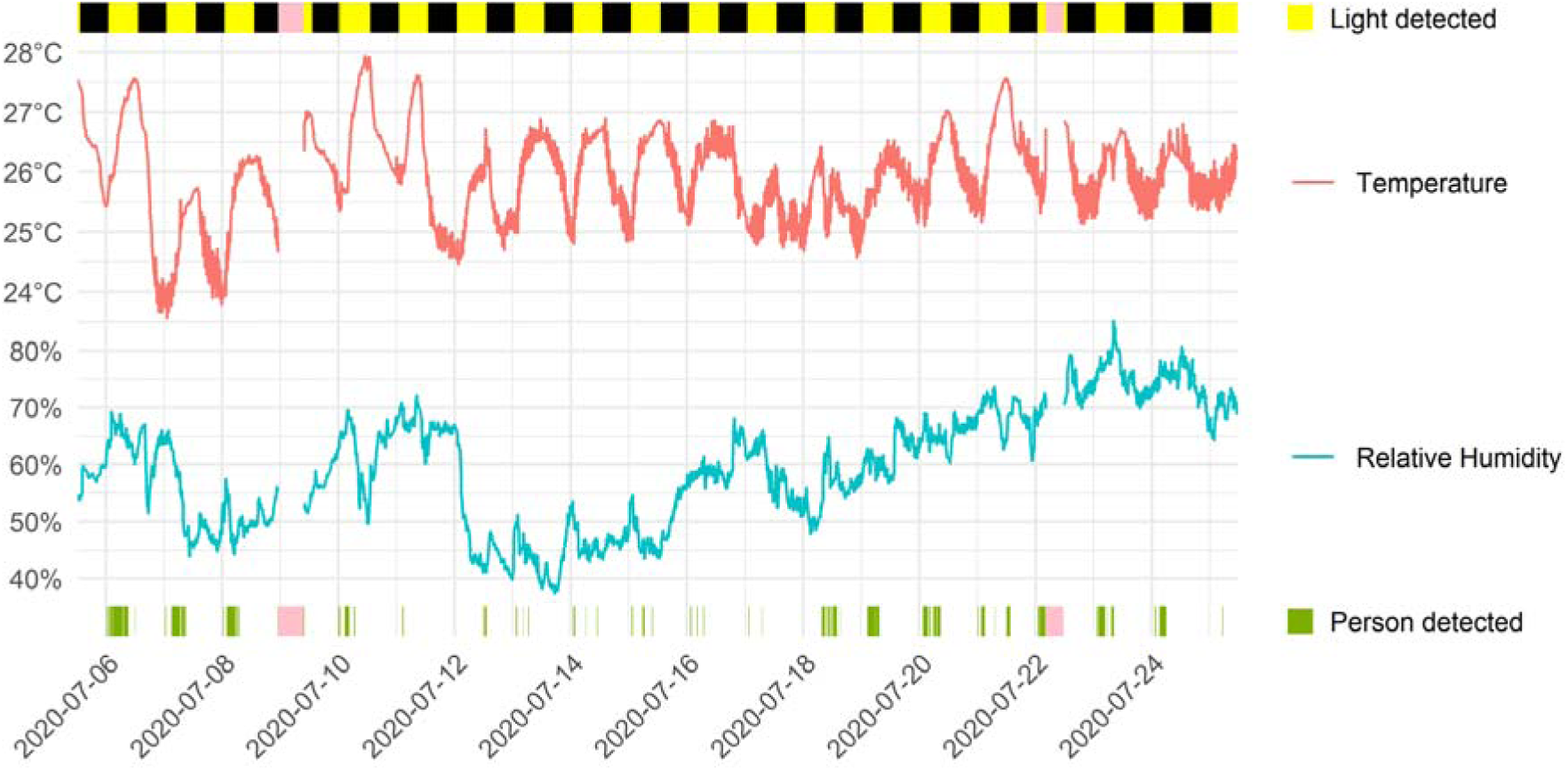
Example of Data Acquired Using MIROSLAV’s Environmental Sensors. *Note.* Pink bars denote missing data.

The mainboard may be extended with a custom board (containing, for example, sensors for specific gases or a microphone for noise level monitoring) through the additional “I2C ENV” connector, which exposes the microcontroller’s I^2^C bus.

MIROSLAV also supports operation as a standalone environmental monitor. This configuration only requires the MIROSLAV mainboard with an attached PIR sensor, and a power supply such as a smartphone charger.

#### Power Boards

The cage sensor arrays and the mainboard may be powered by 5V supplies, commonly found as smartphone chargers. Consideration needs to be given to make sure the power supply’s current capabilities meet a MIROSLAV’s needs, particularly in regard to the number of sensors. However, a MIROSLAV’s sensors may be distributed across multiple power boards, each with its own power supply. If necessary, the power supply boards may also be adapted based on the provided designs.

The input serialisation stack doesn’t require additional power or connection to the power boards. It is always powered directly from the mainboard via the same connection responsible for data transmission.

### Infrastructure

#### Wireless Data Acquisition

To utilise MIROSLAV’s main mode of data acquisition, a Wi-Fi network is required. Data is sent using the MQTT protocol which is designed to support applications with thousands of messages per second exchanged between many senders and receivers (Banks et al., 2019; Mishra et al., 2021), with a single device, e.g. a personal computer (PC) or a router, acting as a “broker”. Multiple on- or off-site computers may be configured to save data from many MIROSLAVs simultaneously for data redundancy, with cloud recording also supported. Additionally, MIROSLAVs are configured by default to synchronise against a local or Internet NTP (Network Time Protocol) server for precise message timestamping across many concurrently running devices.

The default MIROSLAV configuration is set up for 10 messages per second, with each message containing data about the current state of all attached sensor arrays, theoretically allowing for upwards of 100 MIROSLAVs (each with up to 64 cage sensor arrays).

### Direct Data Acquisition via USB Connection

Alternatively, a MIROSLAV may also be attached directly to a computer via USB connection without requiring a wireless network. This mode is limited to one storage device which needs to be placed within the animal habitat in close proximity to the MIROSLAV, reducing robustness. As such, this mode is intended for testing and short-term recording.

### Data acquisition

The accompanying software is used to: i) get current sensor states and transmit them wirelessly to one or more logging PCs; ii) save these readings as compressed raw data; iii) preprocess the raw data for subsequent experiment-dependent analyses. All software is licensed under the GNU Public License version 3 (GPLv3) (Stallman, 2007).

### MIROSLAV Mainboard

The MIROSLAV’s onboard microcontroller is programmed in C++ atop the Arduino software framework. All user-configurable parameters (network settings, polling rates, enabling/disabling environmental monitoring) may be adjusted in the “config.h” file without the need to modify C++/Arduino code. A running MIROSLAV can be updated with a new configuration remotely through a web browser using the *HTTPUpdateServer* library included in the Arduino ESP32 core (Espressif Systems, 2016/2024).

### Data Logging to a PC with Record-a-SLAV

A Python script, *Record-a-SLAV.py*, runs on logging PCs continuously for the duration of the experiment and saves MIROSLAV data to compressed files. The same script can be run on many PCs simultaneously for redundancy.

### Data analysis

Raw data from the MIROSLAV hardware setup is stored as a compressed textual file.

Data analysis steps consist of: i) cleaning and preprocessing, to generate a tall, tidy (Wickham, 2014) columnar file with appropriate labels and discarded faulty data; ii) exploratory data analysis, for initial data examination and validation; iii) statistical data analysis, to test for the existence of putative treatment/intervention effects.

For this process, we have prepared a versatile Python/R-based pipeline, written with the objective of giving the researcher full control over their data and all steps involved. The scripts are highly configurable and intermediate files are generated at multiple steps to allow for combined analysis using existing software libraries and packages such as *Cosinor* (Sachs, 2023), *Kronos* (Bastiaanssen et al., 2023), *GLMMcosinor* (Parsons et al., 2024), and others.

### Data Preprocessing with Prepare-a-SLAV

The Python preprocessing script, *1_Prepare-a-SLAV.py*, utilises the *mirofile* library to load raw MIROSLAV data into a *pandas* data frame. Appropriate animal IDs are applied, and the data is downsampled from the default MIROSLAV sampling rate (10 sensor readings/binary values per second) to an arbitrary, user-defined time interval (bin). For example, resampling a sensor’s readings to a 5-minute bin will turn 864.300 binary values sampled over one day into 288 decimal values by taking the mean of every 3.000 consecutive readings. Narrow bins (seconds, minutes) can give insight into short events, while wide bins (tens of minutes, hours) facilitate visual inspection of hourly, daily, or weekly trends. All parameters are set via a separate, human-readable TOML (*Tom’s Obvious Minimal Language* (Preston-Werner et al., 2021)) configuration file, *1_Prepare-a- SLAV_config.toml* .

### Data Cleaning with TidySLAV

The Python cleaning script, *2_TidySLAV.py*, primarily turns the table into a tidy, tall format (Wickham, 2014). This enables easy addition of experimental metadata and later analysis in a statistical software package. Additionally, some columns are removed, such as disconnected and faulty sensors, as specified by the user. Brief periods when operational sensors were disconnected (e.g. due to behavioural testing or bedding change) are automatically recognised and set to *NA*, indicating an unknown activity state. Finally, the time series are standardised according to each sensor’s respective baseline measurements taken before treatment application.

### Circadian Rhythm Analysis: MIROSine

The objective of *MIROSine* (provided as the R script *3_MIROSine.R*) is to analyse the animals’ circadian rhythm of locomotor activity over the duration of the experiment with the ability to control for confounding variables in a computationally efficient manner.

Simple periodicity with a defined period (24 hours in the case of a circadian rhythm) is well-described by the sinusoidal function:

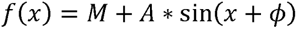

The parameters of the function are i) *M* – the midline, the sinusoid’s equilibrium; ii) *A* – the amplitude, the sinusoid’s maximum offset from its midline; iii) φ – the phase shift, or the horizontal offset of the sinusoid’s peaks and troughs. Cornelissen defines these parameters as the MESOR (Midline Estimating Statistic of Rhythm (MESOR), amplitude, and acrophase, respectively, and proposes the *cosinor* statistical model to estimate them (Cornelissen, 2014). Note that while the terms “acrophase”, “phase shift”, and “peak time of day” refer to the same biological phenomenon, the exact definitions are somewhat different. A visualisation of these parameters is presented in Table 1.

**Table 1.**
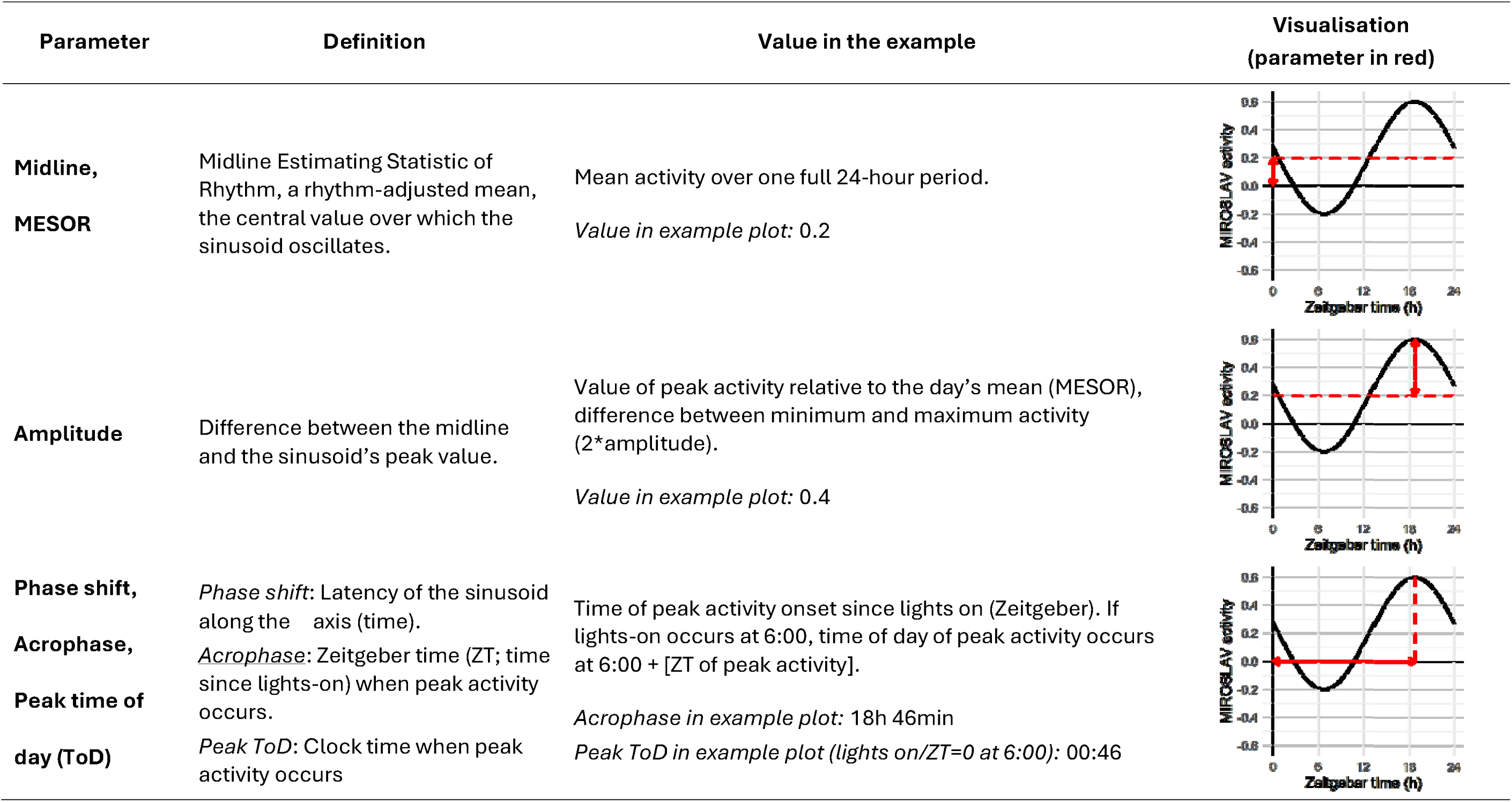
Summary of the Three Sinusoid Parameters.

| Parameter | Definition | Value in the example | Visualisation<br>(parameter in red) |
| --- | --- | --- | --- |
| **Midline,**<br>**MESOR** | Midline Estimating Statistic of Rhythm, a rhythm-adjusted mean, the central value over which the sinusoid oscillates. | Mean activity over one full 24-hour period.<br>*Value in example plot: 0.2* |  |
| **Amplitude** | Difference between the midline and the sinusoid's peak value. | Value of peak activity relative to the day's mean (MESOR), difference between minimum and maximum activity (2*amplitude).<br>*Value in example plot: 0.4* |  |
| **Phase shift,**<br>**Acrophase,**<br>**Peak time of**<br>**day (ToD)** | *Phase shift:* Latency of the sinusoid along the axis (time).<br>*Acrophase:* Zeitgeber time (ZT; time since lights-on) when peak activity occurs.<br>*Peak ToD:* Clock time when peak activity occurs | Time of peak activity onset since lights on (Zeitgeber). If lights-on occurs at 6:00, time of day of peak activity occurs at 6:00 + [ZT of peak activity].<br>*Acrophase in example plot: 18h 46min*<br>*Peak ToD in example plot (lights on/ZT=0 at 6:00): 00:46* |  |

In a manner similar to Cornelissen’s approach, MIROSine estimates these parameters for every day of each sensor’s readings across the duration of the experiment using R’s *lm* function (R Core Team, 2024).

These parameters may then be estimated for an entire group across a specified time interval, as exemplified in the following exploratory and statistical analysis steps performed by MIRO The Explorer and StatistiSLAV, respectively. Care should be taken while interpreting these parameters for days when cage arrays have been disconnected for prolonged periods of time (e.g. due to behavioural testing), as the missing values could skew model fit. TidySLAV heatmaps can be used to find and assess intervals of missing values.

Exploratory Analysis of Circadian Parameters with MIRO The Explorer. Rhythm change is assessed by observing the temporal dynamics of these parameters within and between the experimental groups over the duration of the experiment. MIRO The Explorer offers visual assessment via two graphical approaches by utilising the *ggplot2* package (Wickham, 2016): *Parameter View* and *Rhythm Simulation* . Events such as behavioural testing and bedding changes are also labelled in these visualisations.

Parameter View shows the changes of sinusoid parameters (midline, amplitude, phase shift) per group across the days of the experiment. One time series plot is produced for each parameter, and all groups’ daily means and standard deviations of the respective parameter are shown (PoC study example: Figure 6A).

Rhythm Simulation draws group circadian activity curves throughout the experiment. A group’s daily medians of the three sinusoid parameters are used to reconstruct a group-representative sinusoid for that day. This is repeated for all experimental days, and a smoothed curve is drawn through the simulated values (proof-of-concept study example: Figure 6B).

### Statistical Analysis of Circadian Rhythm Parameters with StatistiSLAV

Statistical analysis is performed using the R script 4_StatistiSLAV.R. Intervals of interest for comparison are selected. One interval may span multiple consecutive days, provided that MESOR, amplitude, and acrophase are stationary across the selected days within each treatment group based on the exploratory analyses.

A generalised linear mixed effects model with restricted maximum likelihood estimation is fitted for each of the three parameters using the glmmTMB package (Brooks et al., 2017). Residuals are simulated using the DHARMa package (Hartig, 2022) and its diagnostic plots are inspected for model assumption violations. Estimated marginal means of the parameters are produced using the emmeans package (Lenth, 2024). The data and model specification were checked against the workflow for linear modelling in R by Santon et al. (Santon et al., 2023).

Two types of visualisations are produced for each time interval. The first are dot plots showing estimated parameter means and 95% confidence intervals for all treatment groups, drawn for each of the sinusoid parameters. The estimated group means of the three parameters are used to reconstruct each group’s sinusoidal representation of circadian activity. The two sinusoids are overlayed in one plot representing each group’s circadian activity for easier interpretation on how the parameter differences affect circadian activity (proof-of-concept study example: Figure 7). This intuitive visualisation is inspired by the cosinoRmixedeffects package developed by Hou et al. (Hou et al., 2021).

Finally, hypothesis testing of group differences per parameter is performed using the emmeans package’s contrasts function. The Holm method is suggested to control the family-wise error rate.

### A Proof-of-Concept Study: Circadian Dysrhythmia in a Rat Model of Sporadic Alzheimer’s Disease

The experiment was performed to establish a comprehensive behavioural phenotype of the streptozotocin-induced rat model of sporadic Alzheimer’s disease (manuscript in preparation). A subset of the data, presented in this work, was also used to demonstrate the validity of the MIROSLAV method, as per the Reduce principle of the 3Rs (Percie du Sert et al., 2020; Russell & Burch, 1992).

### The rat model of sporadic Alzheimer’s disease

The sporadic Alzheimer’s disease (AD) model was induced via intracerebroventricular (icv) injection of streptozotocin (STZ) following a standard procedure used and optimised by our research group (Homolak et al., 2021, 2023; Salkovic-Petrisic et al., 2021) since the model’s introduction by Lacković and Šalković (Lacković & Šalković, 1990), and Hoyer et al. (Mayer et al., 1990). Thirty 3-month-old male Wistar rats, bred at the Department of Pharmacology, University of Zagreb School of Medicine, were randomly divided into a treatment and a control group. The control group (CTR, n = 15) was given 0.05 M citrate buffer (pH 4.5) intracerebroventricularly, and for the streptozotocin-treated group (STZ, n = 15), STZ (1.5 mg/kg) was freshly dissolved in the citrate buffer. Citrate buffer or STZ solution (2 μL) was given into each ventricle twice across 48 hours, at coordinates −1.5 mm posterior, ±1.5 mm lateral, and +4 mm ventral to the bregma (Homolak et al., 2023; Knezovic et al., 2017, 2023; Noble et al., 1967). For the procedure, the animals were anaesthetised with an intraperitoneal injection of ketamine/xylazine (70/7 mg/kg).

### Animal housing

Animals were kept at the Department of Pharmacology’s licensed animal facility (HR-POK-007). The facility was kept at 21-23°C and 40-70% relative humidity. The lighting cycle was 12:12 h, with lights on from 05:46 (Zeitgeber time [ZT] 0) to 17:46 (ZT 12). The animals were single-housed starting 7 days prior to STZ administration. Animals had access to standard food pellets and water ad libitum. Bedding change was performed on a weekly basis. All animal procedures were approved by the Croatian Ministry of Agriculture (EP 378/2022).

### MIROSLAV recording and other in vivo procedures

MIROSLAV sensor arrays were fitted to the cage 4 days prior to STZ administration (day 0) to record baseline activity measurements. Activity was recorded for the following 25 days. Notable events of interest for MIROSLAV recording were weighing, bedding change, and behavioural testing. Animals were weighed on a weekly basis during bedding change.

Behavioural testing was performed from day 16 to day 20 in the following order: i) Open field; ii) Novel object recognition; iii) Social interaction; iv) Morris water maze; v) PASTA startle and prepulse inhibition (Virag et al., 2021). A schematic illustrating the timeline is shown in Figure 5.

**Figure 5.**
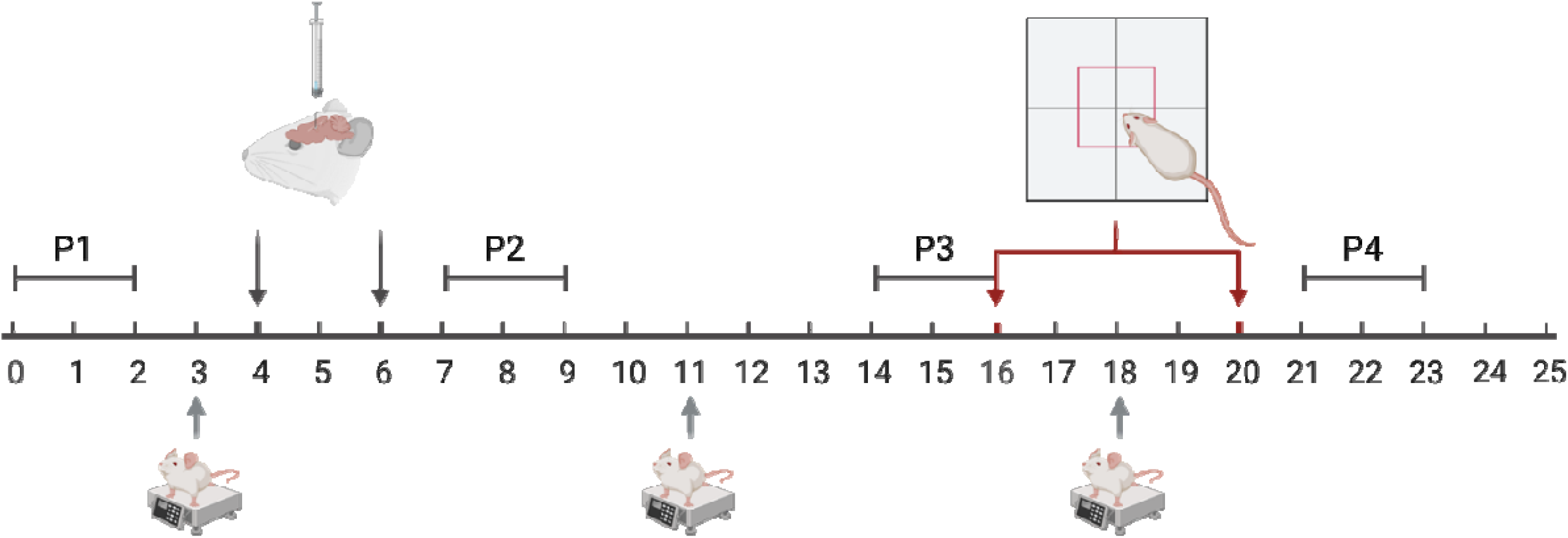
A Schematic of the Experiment Timeline. *Note.* MIROSLAV recording was performed throughout the 25-day period. P1, P2, P3, and P4 denote the four periods taken for statistical analysis with StatistiSLAV. Black arrows signify days when intracerebroventricular application of STZ or citrate buffer was performed. Red arrows signify the period when behaviour testing was performed. Gray arrows signify days of weighing and bedding change.

### Data analysis

Data analysis was performed using the MIROSLAV software suite. Data preparation was performed with Python using Prepare-a-SLAV and TidySLAV. Raw logs were resampled to 1- minute bins using Prepare-a-SLAV. TidySLAV was used to examine the data for technical faults and export it into a tidy, long format.

Circadian activity was analysed using MIROSine to examine the dynamics of 24-hour rhythm amplitude, midline, and phase shift over the course of the experiment. Exploratory data analysis was performed in R using MIRO The Explorer to gain an insight into overall trends and select time periods of particular interest for statistical analysis.

Statistical models were fitted using the R script StatistiSLAV to estimate group MESORs, amplitudes, and phase shifts, as well as respective contrasts between control and STZ treatment groups. Family-wise error rate was controlled using the Holm correction. Alpha was set to 5%.

Full MIRO The Explorer and StatistiSLAV analyses are available in the MIROSLAV analysis repository, which also contains links to live online notebooks (Virag, 2024b).

## Results

Based on MIRO The Explorer analyses (Figure 6), periods of interests for further analysis were selected: i) baseline measurements (days 0 and 1); ii) acute rhythm changes within days of streptozotocin application (days 7 and 8); iii) stabilised rhythms one week after STZ application (days 14 and 15); iv) rhythms after behavioural testing (days 21 and 22) (Figure 5).

**Figure 6.**
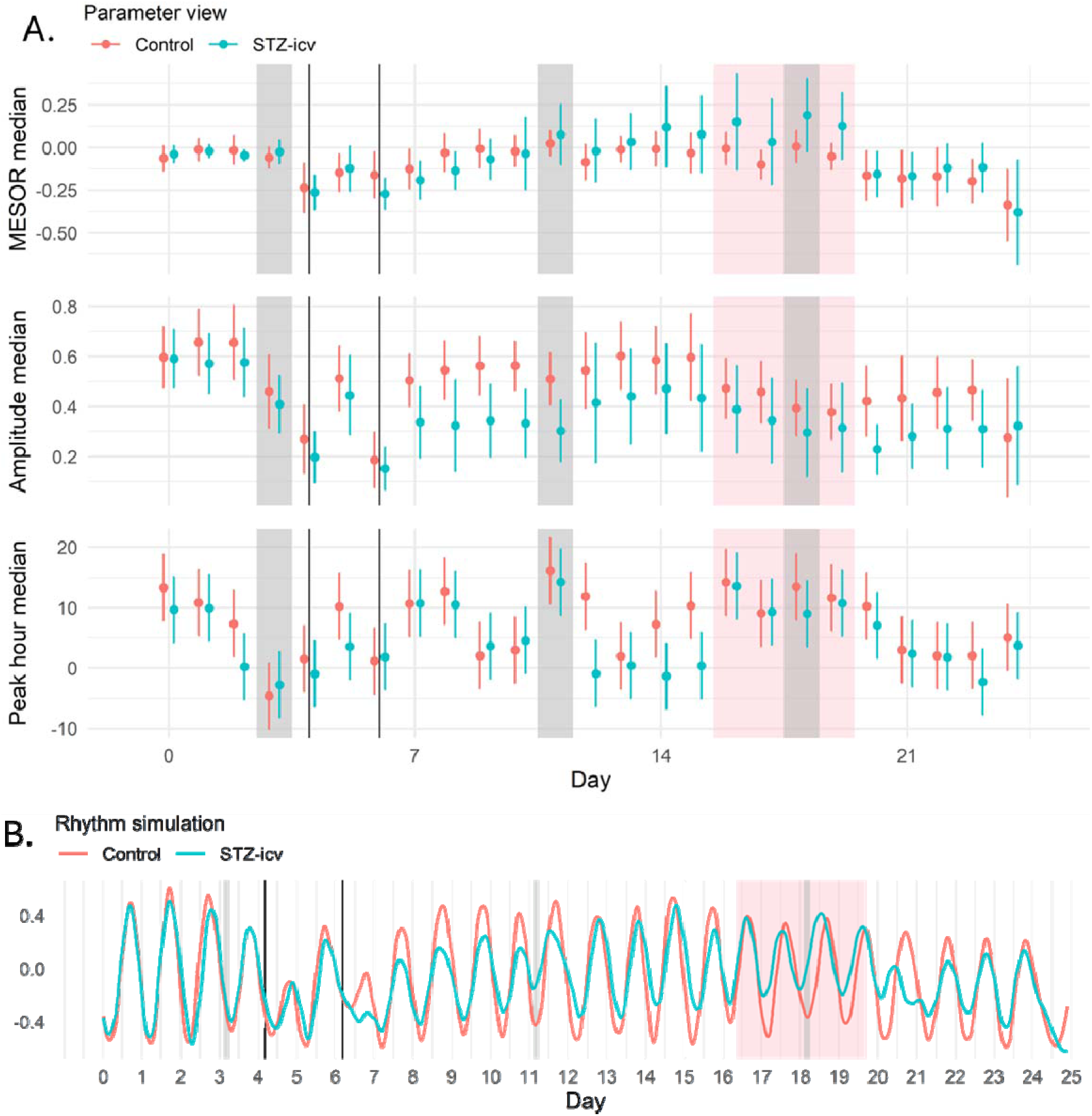
Proof-of-Concept Study Exploratory Analysis Plots Produced with MIRO The Explorer. *Note.* Gray bars signify the days/times of animal weighing and cage bedding change. Gray vertical lines signify starts of each labelled 24-hour period. Black vertical lines at days 4 and 6 signify intracerebroventricular injections. Behavioural testing was performed during the period highlighted in red. Panel A: Parameter view shows temporal dynamics of the three sinusoid parameters. Panel B: Rhythm simulation reconstructs group-representative activity, calculated from group medians of the three sinusoid parameters for each day.

MIROSLAV measurements before streptozotocin administration show no difference between animals at baseline (Figure 7A). Within the two days following model induction, a lower nocturnal activity peak (zenith) is seen in the STZ-icv group (Figure 7B). One week later, the zenith is reestablished, but the diurnal activity trough (nadir) is higher in the STZ-icv group. STZ-icv-treated animals also exhibit a latency in the zenith onset of approximately 50 minutes at this timepoint (Figure 7C). During behavioural testing, MIRO The Explorer indicates a higher diurnal activity in the STZ-icv group, as well as a lingering circadian disruption in the following days (Figure 6). StatistiSLAV analysis on the second and third day after behavioural testing shows a reduction in amplitude, indicating both a lower nocturnal zenith, and a higher diurnal nadir in the STZ-icv group (Figure 7D).

**Figure 7.**
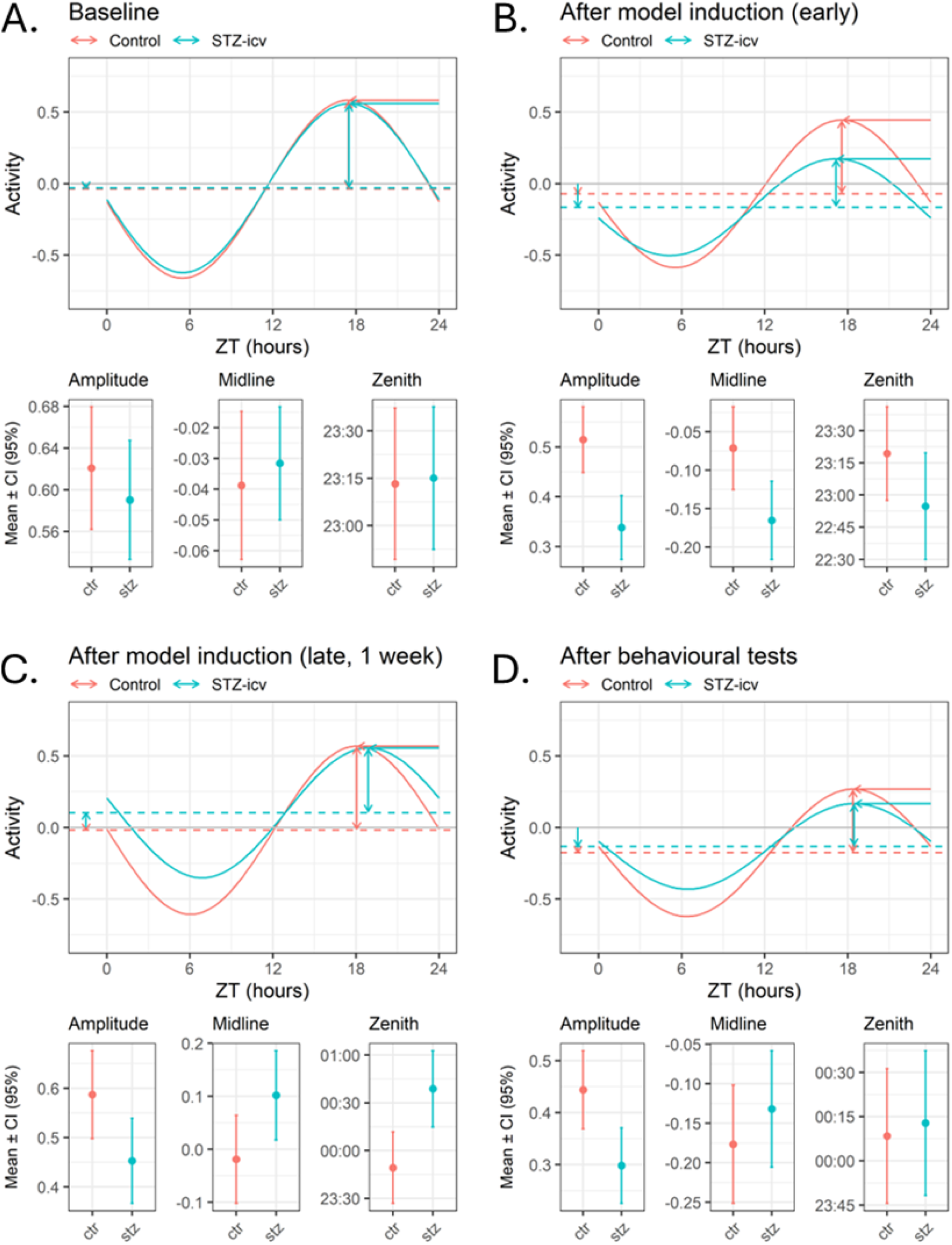
Proof-of-Concept Study Statistical Analysis Plots Produced with StatistiSLAV. *Note.* The figure shows group estimates of the three sinusoid parameters reconstructed sinusoidal representations at four timepoints of interest.

## Discussion

Due to the general limitations of proprietary, commercial home cage monitoring systems, and by building on pioneering ideas of open source tools such as RAD (Matikainen-Ankney et al., 2019) and COMPASS (Brown et al., 2016), we have decided to develop MIROSLAV as a modular platform with the goal of supporting as many experimental settings as possible. To achieve the necessary robustness, over the years of prototype development and testing, we have identified common points of failure and implemented this experience in our design to support high-throughput recording of rodent locomotor activity in large studies.

MIROSLAV has shown that its design considerations resulted in a platform with real- word utility by allowing us to non-invasively assess and describe the circadian dysrhythmia in the rat sporadic Alzheimer’s disease model induced by intracerebroventricular streptozotocin, as well as to evaluate the effect of behavioural testing on the animals. We present these results thoroughly processed as part of the proof-of-concept study here, but MIROSLAV’s earlier prototypes have shown preliminary findings of circadian dysrhythmia, to the best of our knowledge, for the first time in this model (Virag, 2021), where locomotor disruption has been described in conventional behavioural tests ever since the model’s first behavioural phenotyping by Mayer et al. (Mayer et al., 1990). Following these MIROSLAV results, an extensive behavioural characterisation of the STZ-icv sporadic AD model yielded interesting findings in the context of a link between AD and attention-deficit hyperactivity disorder (Grünblatt et al., 2023), prompting a new avenue of research. This brief example illustrates further benefits that widely available HCM tools could carry.

### What makes MIROSLAV a friend indeed?

#### Hardware and infrastructure design

Addressed hardware and infrastructural considerations making MIROSLAV appropriate for long-term monitoring at scale are listed in Table 2.

**Table 2.** MIROSLAV Hardware Design Choices for Practicality and Robustness.

| Point of failure | Design choices | Impact in daily use |
| --- | --- | --- |
| Cage-mounted sensor array:<br>• Disconnecting the sensor array to remove a cage from the rack<br>• Removing the sensor array from the cage lid | • Sensor array connects to the main unit with a zero-force ethernet/telephone-like connector (RJ9), similar to COMPASS (Brown et al., 2016)<br>• No tools required to fully remove the sensor array from the cage lid | The connector requires grabbing the head and pressing a pin to release the connection, eliminating the need for applying force to the connector and cable-tugging. This approach maintains and prolongs the integrity of the cable over many connect-disconnect cycles. |
| Partial (spurious readings) or complete failure (data loss) of a sensor unit | Two sensor units in each array, sharing one connection to the main unit. | In case of complete failure, data is still recorded by the second sensor. In case of partial failure, TidySLAV delta heatmaps show a high discrepancy between the two sensors' readings, alerting the user. |
| Power outage in the habitat | Main data acquisition unit is based around a microcontroller (as opposed to PC, Raspberry Pi) | Lacking a filesystem and operating system, microcontrollers do not suffer from program corruption on power loss, long start-up times, or require reconfiguration.<br>After power is reestablished, data acquisition is resumed within seconds. |
| Disruptive noise in the habitat, physical damage to the device from habitat conditions | The WiFi-enabled microcontroller stores no data in the habitat, but transmits it via WiFi. Requires WiFi coverage of the habitat area (but not Internet access). | A separate device, placed outside of the habitat, but connected to the same network, accepts MIROSLAV data and records it permanently. In case of physical damage to the MIROSLAV device itself, all components can be immediately replaced without additional configuration, ensuring minimal loss of data. |

| Point of failure | Design choices | Result |
| --- | --- | --- |
| Failure of a recording device | Wireless data transmission via the MQTT protocol, which allows dynamic connection of recording devices | An MQTT “broker” forwards data to as many on- or off-site recording devices as needed, all of which retain copies of the data. Devices may connect (“subscribe”) or disconnect dynamically without disrupting the data acquisition process. |
| Time synchronisation of sensor readings from multiple operating devices | Network Time Protocol (NTP) synchronisation of all devices, timestamping on message send and receive. | All MIROSLAV and recording devices synchronise their clocks against the same NTP server (which can be deployed on a local network without requiring Internet access) and record the exact date and time when the message was both sent (MIROSLAV) and received (recording device). |

*Note.* This table describes common home cage monitoring-related failures discovered in empirical testing, MIROSLAV design choices made to mitigate or eliminate points of failure, and how the choices translate to device robustness in normal MIROSLAV operation.

Our experience has shown the system to be robust used as four parallelised MIROSLAV units simultaneously recording activity from 87 cages to two recording computers over the same wireless network, and a third one off-site via a secure Internet connection. Having multiple wirelessly connected recording devices outside of the animal habitats was of great practical value in our experiments – it enabled remote animal and equipment checks (in addition to regular, in-person daily checks), without increasing risk of contamination as no habitat entry was required. Considering the lack of issues in a large-scale deployment and high theoretical limits of operation (described above in the Infrastructure section of the MIROSLAV device description), we are confident the MIROSLAV platform could be used in deployments of hundreds of cage units. Finally, of note is the low cost of the hardware system, amounting to under 1,500 EUR for a full package needed to build five MIROSLAVs with a total of 120 cage sensor arrays and a large quantity of spare parts in case of damage. A full bill of materials available in the MIROSLAV hardware repository (Virag, 2024a).

#### Software design and data quality

While the utility of continuously operating HCM devices in virtually all types of experimental designs is indisputable, there are many practical considerations when working with the acquired data. Today, the industry and the scientific community are hard at work developing new techniques handling the immense quantity of data these types of devices produce, in terms of storage, processing, and ensuring data quality (Baran et al., 2021; D’Argenio, 2018; Fuochi et al., 2024).

These concerns apply to MIROSLAV as well – after obtaining raw data, it needs to be analysed to garner biological insight. The MIROSLAV software package was developed to transparently guide the researcher through three principal steps of the data analysis process, as described in Table 3.

**Table 3.**
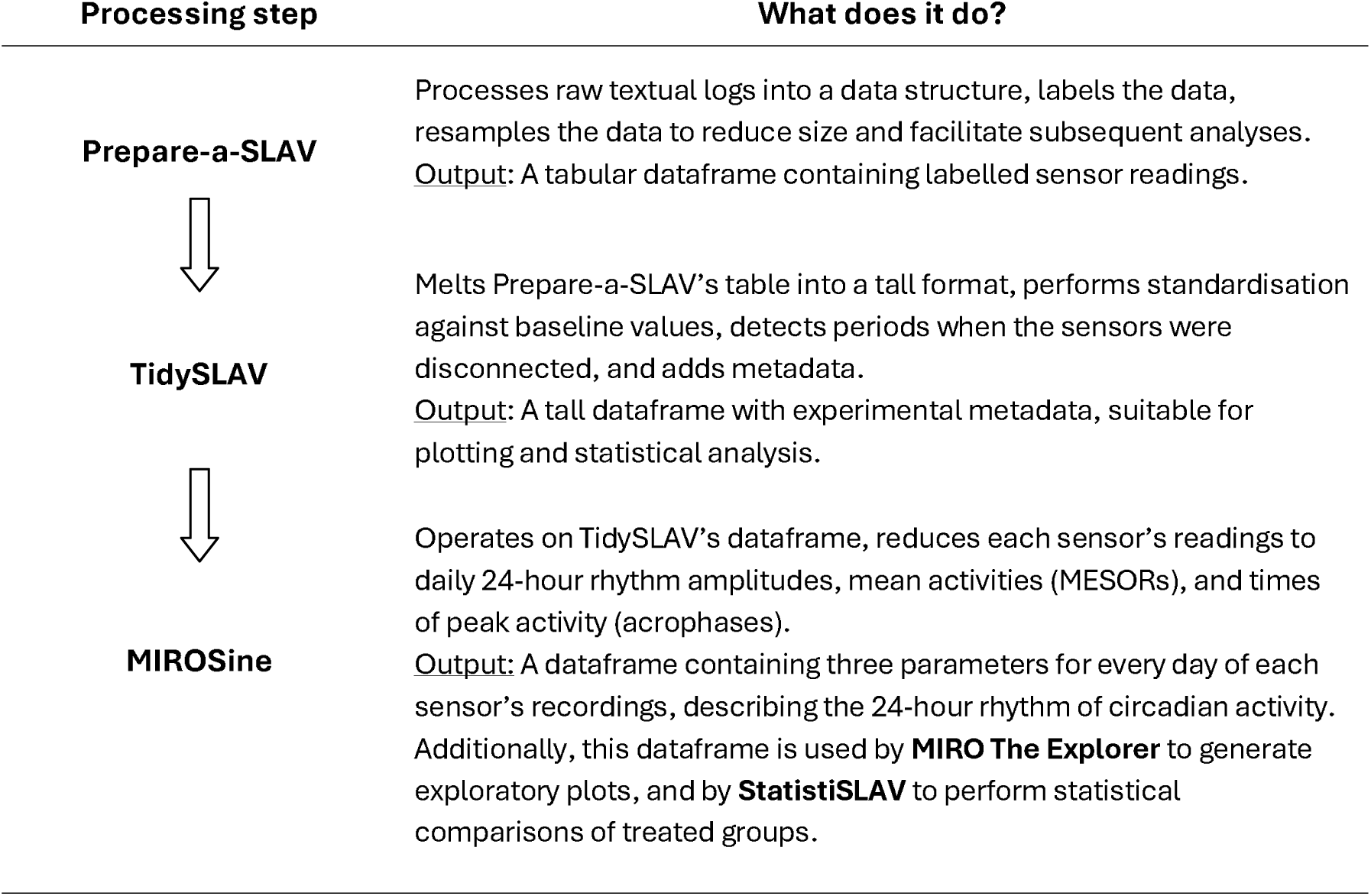
Description of the MIROSLAV Software Toolkit Workflow.

| Processing step | What does it do? |
| --- | --- |
| **Prepare-a-SLAV** | Processes raw textual logs into a data structure, labels the data, resamples the data to reduce size and facilitate subsequent analyses.<br><u>Output:</u> A tabular dataframe containing labelled sensor readings. |
| **TidySLAV** | Melts Prepare-a-SLAV's table into a tall format, performs standardisation against baseline values, detects periods when the sensors were disconnected, and adds metadata.<br><u>Output:</u> A tall dataframe with experimental metadata, suitable for plotting and statistical analysis. |
| **MIROSine** | Operates on TidySLAV's dataframe, reduces each sensor's readings to daily 24-hour rhythm amplitudes, mean activities (MESORs), and times of peak activity (acrophases).<br><u>Output:</u> A dataframe containing three parameters for every day of each sensor's recordings, describing the 24-hour rhythm of circadian activity. Additionally, this dataframe is used by **MIRO The Explorer** to generate exploratory plots, and by **StatistiSLAV** to perform statistical comparisons of treated groups. |

In the MIROSLAV software suite, we would like to highlight MIROSine as a novel approach to analysing locomotor activity data with a focus on the animals’ circadian rhythm. Without assuming static circadian rhythm parameters over the course of an experiment, the approach allows for: i) intuitive visualisations of circadian rhythm dynamics over time; ii) the application of mixed regression models to the data for statistical analysis, with the inclusion of appropriate covariates, interactions, and random effects. Unlike bootstrapping approaches, it is not computationally intensive, with MIROSine analyses with our example data being successfully completed within minutes on a personal laptop.

#### MIROSLAV as an open, modular platform

The above considerations led to a reliable method which has so far found extensive use in our laboratory. However, to achieve high configurability as a prerequisite for supporting possible future experimental designs and wider use outside of our own context, MIROSLAV needed to be designed as an open and modular platform (Gavras & Kostakis, 2021).

For example, a MIROSLAV unit doesn’t need to be used as a home cage monitoring tool. A standalone mainboard can function as a small, Wi-Fi-connected monitor of habitat environmental parameters. Even though one unit supports up to 64 cages, the modular input stack can be reduced to a single board, e.g. to spread out many cages across multiple MIROSLAV units as a means of reducing the number of failure points. This option is also feasible due to MIROSLAV’s low cost. Furthermore, to change the scale from a smaller, preliminary experiment to a large-scale one, or vice versa, the power supply is perhaps a less obvious consideration, but one of critical importance. As large 5V power supplies are somewhat difficult to source due to availability and price, the load of a larger number of sensors can be distributed across a larger number of standard, smaller power supplies which are trivial to add. This architecture can be used as a foundation for other novel tools, and we are currently developing a home cage conditioning apparatus based on the MIROSLAV architecture in our laboratory (Virag et al., 2024).

From the software side, we recognise that freely available code in a repository is insufficient for replicability and reuse. MIROSLAV analysis code is written as notebooks (Grayson et al., 2022; Ivimey-Cook et al., 2023; Paxton, 2018/2023; Pimentel et al., 2021) – documents with human-readable, formatted text serving as documentation, with interspersed code cells where configuration parameters can be modified and code can be executed to produce the desired results within the document itself. Unlike a graphical user interface, this approach allows researchers to quickly become intimately familiar with how their data is handled throughout the entire process. As the notebook is mainly composed of formatted text describing the code cells’ processes and configuration, a researcher with no knowledge of programming can simply adjust the available parameters to adapt the analysis for their needs. Similarly, the extensive code descriptions required by this approach facilitate contributions by researchers adept at programming, who can submit their code changes to the repository and continue to build up the MIROSLAV platform through a collaborative effort seen in many open source projects, from DeepLabCut (Mathis et al., 2018) to the Linux kernel itself (Moon & Sproull, 2000). Finally, the analysis code is also modularised according to the steps outlined previously (Table 3). Prepare-a-SLAV, TidySLAV, and the MIROSine tools are completely independent and store a dataframe with a defined format, giving researchers the freedom to seamlessly intercalate other tools at various points in the process.

By designing MIROSLAV to be transparent and highly adaptable at every level, we hope every researcher in need will find a friend indeed in MIROSLAV.

## Limitations

The MIROSLAV system has some inherent limitations.

While general benefits of passive infrared sensors include non-invasiveness and low cost, they aren’t able to discern individual animals in group-housed experimental designs – any movement is registered, regardless of which animal produced it. Furthermore, the HC-SR501 sensor modules used in the construction of MIROSLAV have individual variability, requiring baseline measurements and subsequent standardisation (performed by TidySLAV) to detect deviations from baseline measurements taken with one sensor after an event of interest has occurred (e.g. application of pharmacotherapy). The latter limitation can be mitigated, allowing direct and immediate comparison of readings from different sensor arrays by fitting the MIROSLAV sensor arrays with factory-calibrated sensor modules (e.g. Panasonic EW AMN32111 used by the COMPASS system (Brown et al., 2016)). Their unit price is, however, approximately 10x higher than that of the default HC-SR501 modules.

While PIR sensors are suitable for tracking patterns of overall locomotor activity, due to the nature of the technology, only information on the existence and duration of any movement within a cage can be recorded. To accurately assess the nature of the movement, combining the MIROSLAV system with other methods (e.g. video recording) is advised. The main unit is fitted with unused connectors which can be used to precisely synchronise MIROSLAV devices with other recording devices using hardware synchronisation/triggering platforms such as the open-source Behaviour-recording Device Synchroniser (BeDSy) (Virag et al., 2023/2024).

Furthermore, while all parts of a MIROSLAV device can withstand 70% ethanol or isopropanol and gentle scrubbing, it was not designed to be sterilised with more aggressive solvents or high heat. Although not yet attempted, we expect vaporised hydrogen peroxide sterilisation can be employed without damaging the device (Loo et al., 2010).

### Future development

There are three main objectives in our future work on MIROSLAV:

1. Assessing approaches to overcoming the aforementioned limitations,
2. While MIROSLAV enables real-time access to recorded data, a notification system is planned to alert researchers of changes in animal state or environmental conditions,
3. Work on the documentation will persist as a continuous effort to make MIROSLAV even more approachable by potential users.

## Conclusion

MIROSLAV is a highly modular open-source hardware and software platform for monitoring home cage activity and habitat conditions, capable of non-invasive, high-throughput operation over extended periods in many experimental settings. The comprehensive, extensively documented software toolkit gives the researcher full insight into every step of the data’s journey, from acquisition to statistical analysis, as an advance from proprietary, commercial tools. It is our hope that this adaptable, transparent, and low-cost solution will help many laboratories maximise the data output of their ongoing research projects and entice inter-group collaboration for further development of the open-source ecosystem of tools made by researchers, for researchers. In this spirit, readers are encouraged to contact the authors should they want to build, adapt, or contribute to the MIROSLAV platform.

## Acknowledgements

The authors would like to thank Pia Kahnau and Lars Lewejohann for their valuable feedback on the idea that was subsequently developed into the MIROSine analysis approach during Davor Virag’s visit to their laboratory (Short-Term Scientific Mission funded by the COST- TEATIME [CA20135] project).

The authors would also like to thank laboratory technician Božica Hržan for supervising MIROSLAV devices and ensuring their continuous operation throughout all MIROSLAV experiments, as well as animal caretakers Miroslava Jerković, Tomislav Petanjek, and Anita Šantak for their patience while handling MIROSLAV-equipped cages.

## Declarations

### Ethics Approval

All animal procedures were conducted in accordance with the Croatian Animal Protection Act (Class: 011-01/17-01/77, Reg no: 71-06-0/1-17-2), aligned with European Union legislation. All animal procedures were approved by the Croatian Ministry of Agriculture (EP 378/2022).

### Consent to Participate

Not applicable

### Consent for Publication

Not applicable

### Availability of Data and Materials

All data presented in the study can be found in the MIROSLAV-analysis Zenodo repository: https://doi.org/10.5281/zenodo.12191589. The repository contains raw sensor output from the PoC study, sample environmental data, and the MIROSLAV analysis toolkit used for processing these datasets. As the toolkit is modularised into six tools used sequentially (each tool further processes the output of the previous one), all intermediate datasets are also included, as well as the final analyses presented in this manuscript.

MIROSLAV hardware designs and a full bill of materials (BOM) needed to construct a MIROSLAV package (five MIROSLAVs with a total of 120 cage sensor arrays and a large quantity of spare parts in case of damage) can be found in the MIROSLAV-hardware Zenodo repository: https://doi.org/10.5281/zenodo.12675452.

### Code Availability

All MIROSLAV code is placed in two Zenodo repositories, MIROSLAV-firmware and MIROSLAV-analysis. Continuous development is done on the corresponding GitHub repositories, which the Zenodo repositories reference. Zenodo and GitHub repositories are linked – with each new GitHub release, a snapshot of the entire repository is taken and stored by Zenodo, which also retains repository snapshots of previous versions. Each version contains a note detailing newly introduced changes and updates.

### MIROSLAV Firmware

**Repository URLs.**

- Zenodo DOI: https://doi.org/10.5281/zenodo.12675471
- GitHub URL: https://github.com/davorvr/MIROSLAV-firmware

**Repository Contents.**

Arduino firmware for the MIROSLAV device (MIROSLAVino), as well as data acquisition software (Record-a-SLAV) for PC recording devices.

- MIROSLAV Arduino firmware (MIROSLAVino): https://github.com/davorvr/MIROSLAV-firmware/tree/main/miroslavino
- Data acquisition tool (Record-a-SLAV): https://github.com/davorvr/MIROSLAV-firmware/tree/main/Record-a-SLAV

### MIROSLAV Analysis

**Repository URLs.**

- Zenodo DOI: https://doi.org/10.5281/zenodo.12191589
- GitHub URL: https://github.com/davorvr/MIROSLAV-analysis

**Repository Contents.**

Contains raw data for the PoC study, sample environmental data, and the MIROSLAV analysis toolkit. MIROSLAV home cage data analysis tools are provided in the main directory of the repository as:

- Pure Python or R scripts (file extension: .py or .R)
- Interactive Jupyter Notebooks (file extension: .ipynb)
- Live online Google Colab notebooks (URLs in the repository README file: https://github.com/davorvr/MIROSLAV-analysis/blob/main/README.md).

Environmental data parsing and plotting tools are included in a subdirectory. The repository also contains intermediate and final analysis outputs of the PoC study data from all of the listed tools.

**Components List.**

1. Raw data: https://github.com/davorvr/MIROSLAV-analysis/tree/main/0_raw_logs
2. Data analysis tools (https://github.com/davorvr/MIROSLAV-analysis):

a. Prepare-a-SLAV (data preparation)
b. TidySLAV (data cleaning)
c. MIROSine (24-hour rhythm analysis)
d. MIROSine-based analyses:

i. MIRO The Explorer (exploratory data analysis)
ii. StatistiSLAV (statistical data analysis)
3. EnviroSLAV (environmental monitoring parsing and plotting tools) https://github.com/davorvr/MIROSLAV-analysis/tree/main/E_EnviroSLAV

## Authors’ Contributions

**DV**, **JH**, and **IK** conceptualised the MIROSLAV platform. **DV** developed prototype and final iterations of MIROSLAV electronics, firmware, and infrastructure. **JH** designed and built three iterations of MIROSLAV prototype units. **AMV** designed the final MIROSLAV device, and built it with **DV**, **JH**, **IK**, and **ABP** .

All prototype analysis code was written by **IK**, upon which **DV** wrote the mirofile Python library, Prepare-a-SLAV, and TidySLAV. Exploratory visualisations were conceptualised by **IK** and developed into TidySLAV diagnostic plots and MIRO The Explorer by **DV**, **JH**, **PM**, and **MŠP** . **DV** conceptualised the MIROSine analysis approach and developed it with **JH** and **PŠM**, with critical input from **MC** and **VT**. **DV**, **JH**, and **VT** conceptualised the StatistiSLAV analysis with critical input from **MC** . The MIROSLAV logo used in the repositories was drawn by **PŠM** .

In the PoC experiment, **DV**, **JH**, **ABP**, **AK**, and **JOB** carried out the drug treatments, and **JH** performed the behavioural testing.

**DV** wrote the manuscript. **JH**, **IK**, **AMV**, **ABP**, **PM**, **PŠM**, **AK**, **JOB**, **MC**, **VT**, and **MŠP** commented on the manuscript and provided critical feedback. **MŠP**, mentor of DV and the laboratory’s PI, provided funding and supervised the project.

Figures 1 and 4 were created with BioRender.com.

## Open Practices Statement

A version of this paper was preprinted on bioRxiv: https://doi.org/10.1101/2024.06.25.600592

Experiments were not preregistered.

All MIROSLAV data, from raw data to statistical analysis output, are placed the MIROSLAV-analysis GitHub repository: https://github.com/davorvr/MIROSLAV-analysis This repository is linked with a Zenodo entry: https://doi.org/10.5281/zenodo.12191589.

All MIROSLAV code and hardware designs used in the study are placed in three GitHub repositories:

- MIROSLAV-hardware (Zenodo DOI: https://doi.org/10.5281/zenodo.12675452; GitHub URL: https://github.com/davorvr/MIROSLAV-hardware)
- MIROSLAV-firmware (Zenodo DOI: https://doi.org/10.5281/zenodo.12675471; GitHub URL: https://github.com/davorvr/MIROSLAV-firmware)
- MIROSLAV-analysis (Zenodo DOI: https://doi.org/10.5281/zenodo.12191589; GitHub URL: https://github.com/davorvr/MIROSLAV-analysis)

Furthermore, the analysis code is presented as Jupyter notebooks which can be loaded in the web browser (using Google Colab) via a button on the repositories’ web pages. The notebooks contain documentation and code output, which can be regenerated/replicated in the web browser by rerunning the code in Colab.

Continuous development is done on GitHub. Zenodo repositories are linked, such that each new GitHub release triggers the generation of a new Zenodo snapshot of the entire repository (including all study data). This way, Zenodo DOIs give readers access to both the most recent developments, as well as the full history of the MIROSLAV project, including its state at the time of publications.

All code is licensed with GPLv3, supporting further code sharing and collaboration on the MIROSLAV platform.

